## Supplementary figures and images for "Occurrence and stability of hetero-hexamer associations formed by β-carboxysome CcmK shell components"

### Supplemental Figure 1

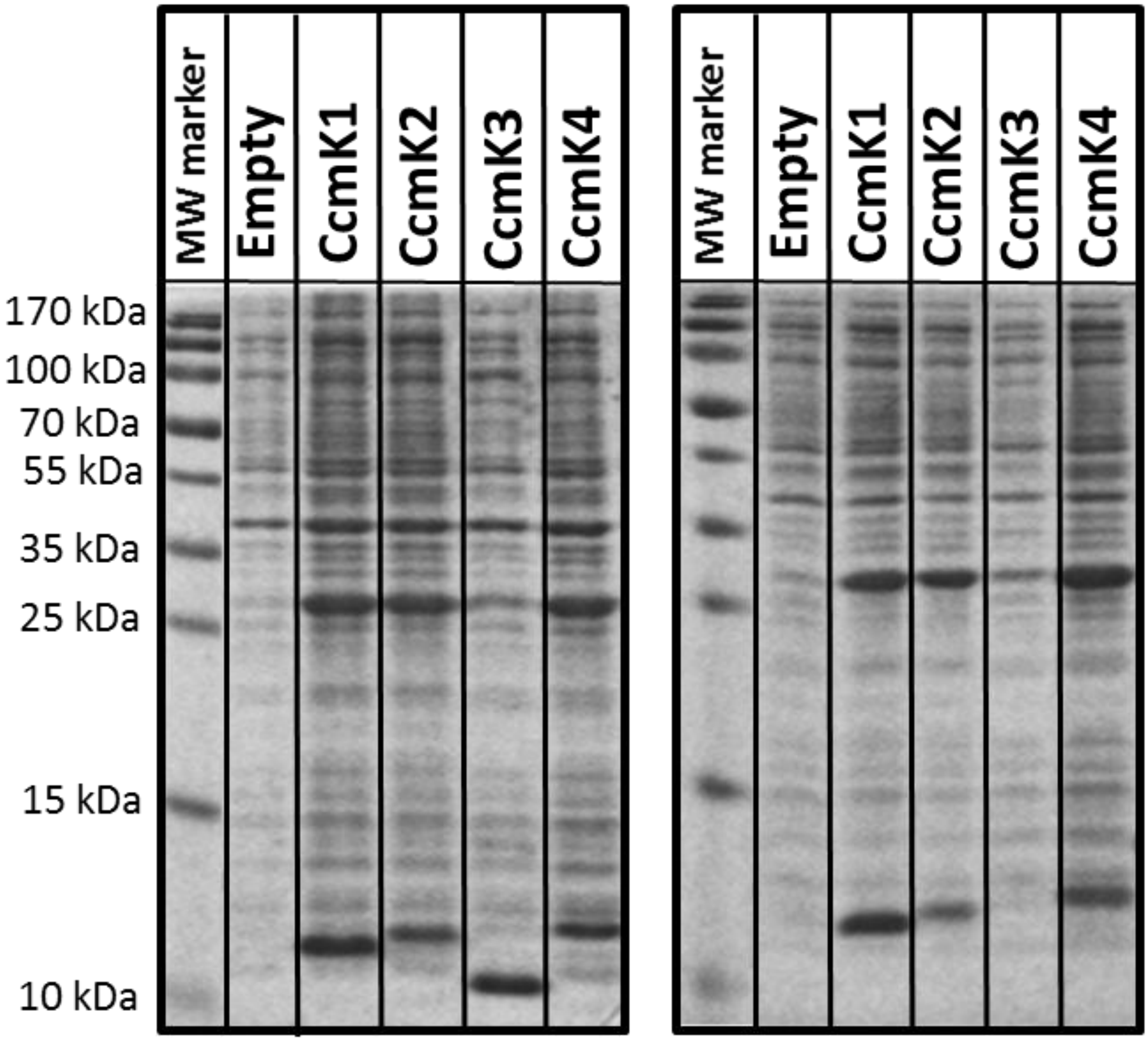

### Supplemental Figure 2

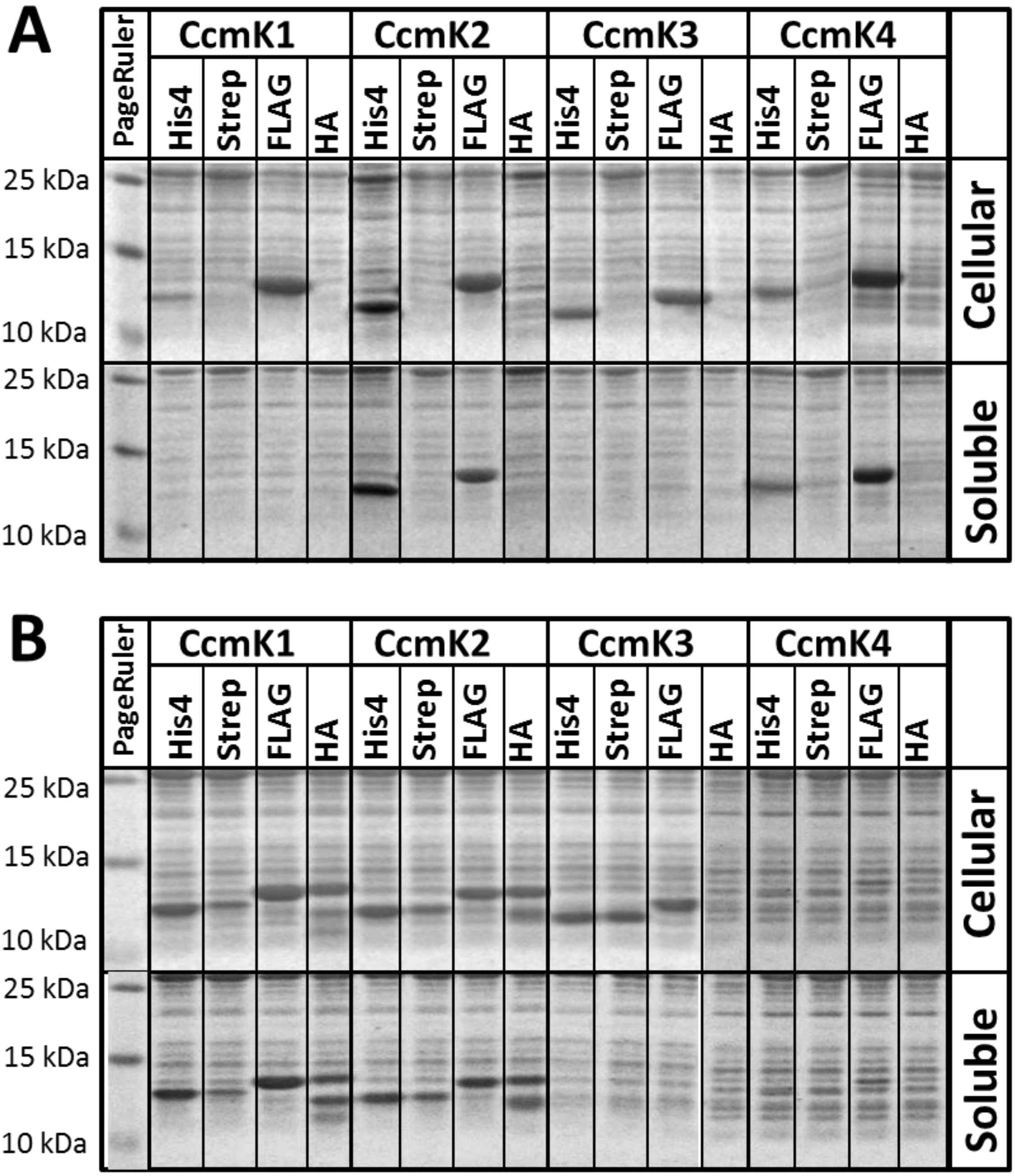

### Supplemental Figure 3

#### Figure S3

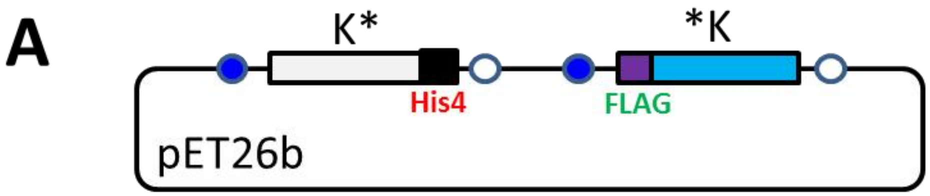


| | | | | |
|---|---|---|---|---|
| **K1*/K1*** | K2*/K1* | K3*/K1* | K4*/K1* | *K4/K1* |
| K1*/K2* | **K2*/K2*** | K3*/K2* | K4*/K2* | *K4/K2* |
| K1*/*K3 | K2*/*K3 | **K3*/*K3** | K4*/*K3 | *K4/*K3 |
| K1*/K3 | K2*/K3 | **K3*/K3** | K4*/K3 | *K4/K3 |
| K1*/K3* | K2*/K3* | **K3*/K3*** | K4*/K3* | *K4/K3* |
| K1*/K4* | K2*/K4* | K3*/K4* | **K4*/K4*** | ***K4/K4*** |

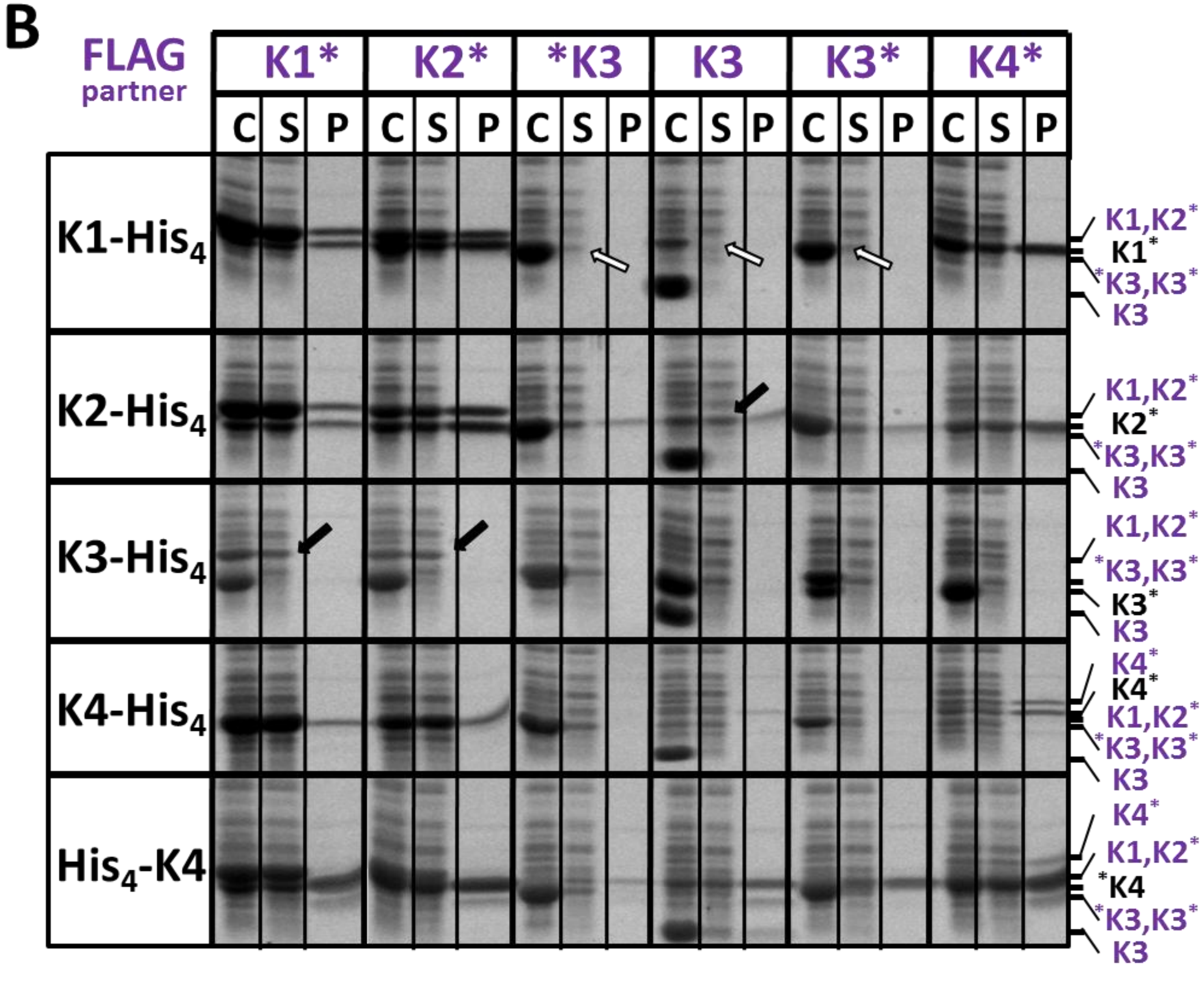

### Supplemental Figure 4

#### Figure S4

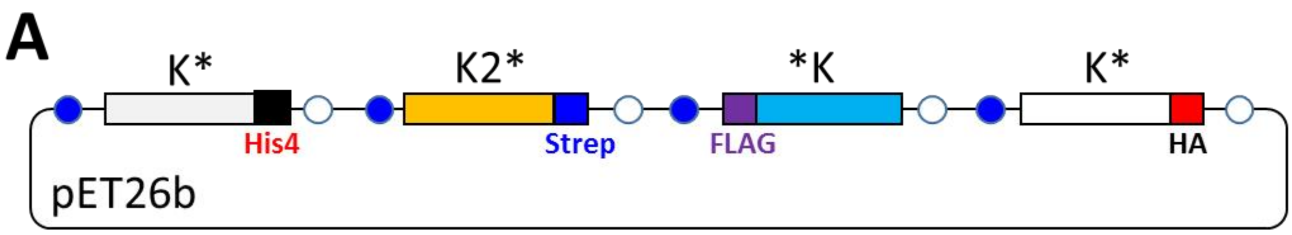


| | |
|---|---|
| K1*/K2*/*K3/K4* | K1*/K2*/K3*/K4* |
| K3*/K2*/*K4/K1* | K3*/K2*/K4*/K1* |
| K4*/K2*/*K3/K1* | K4*/K2*/K3*/K1* |
| *K4/K2*/*K3/K1* | *K4/K2*/K3*/K1* |

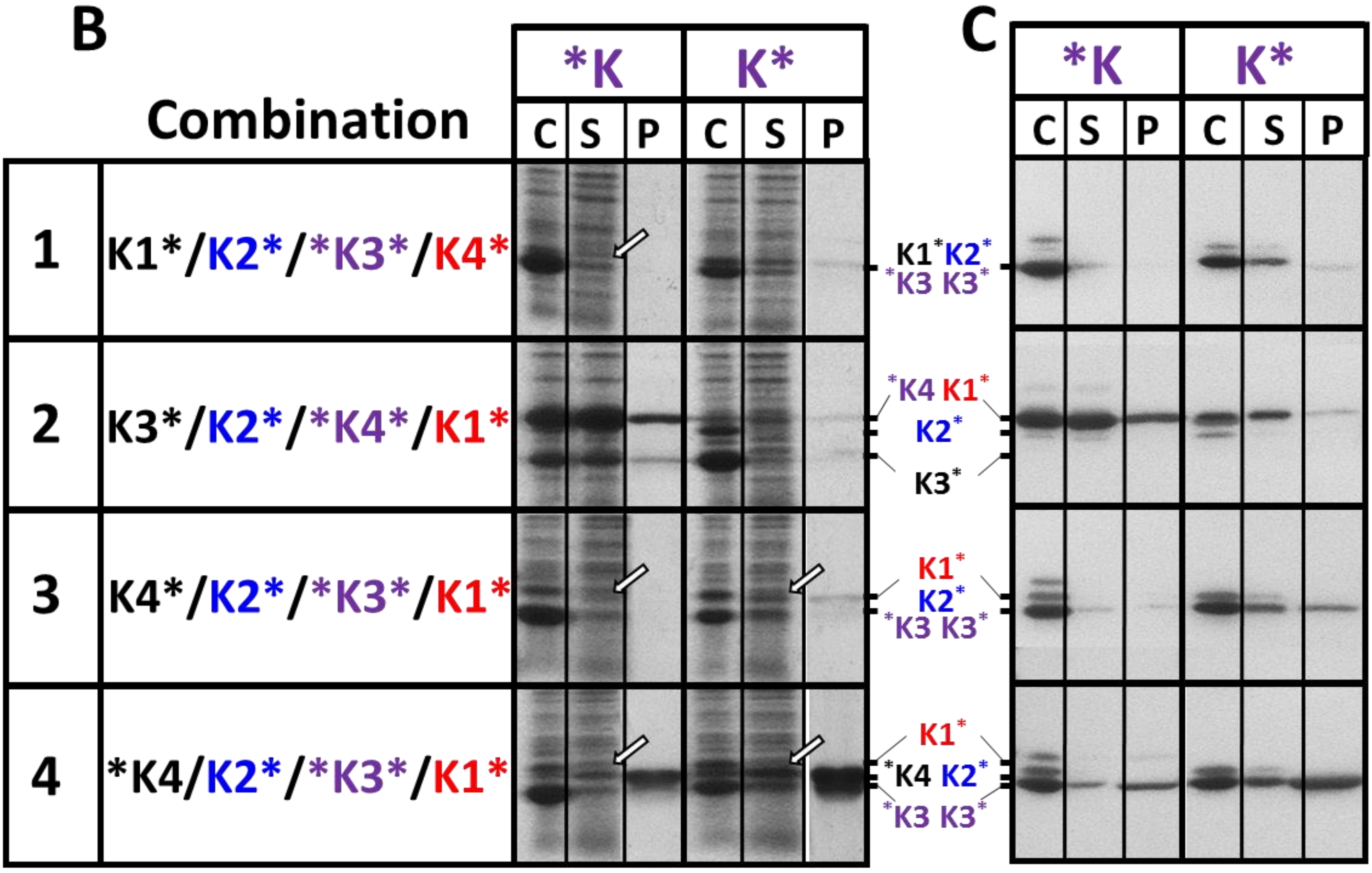

### Supplemental Figure 5

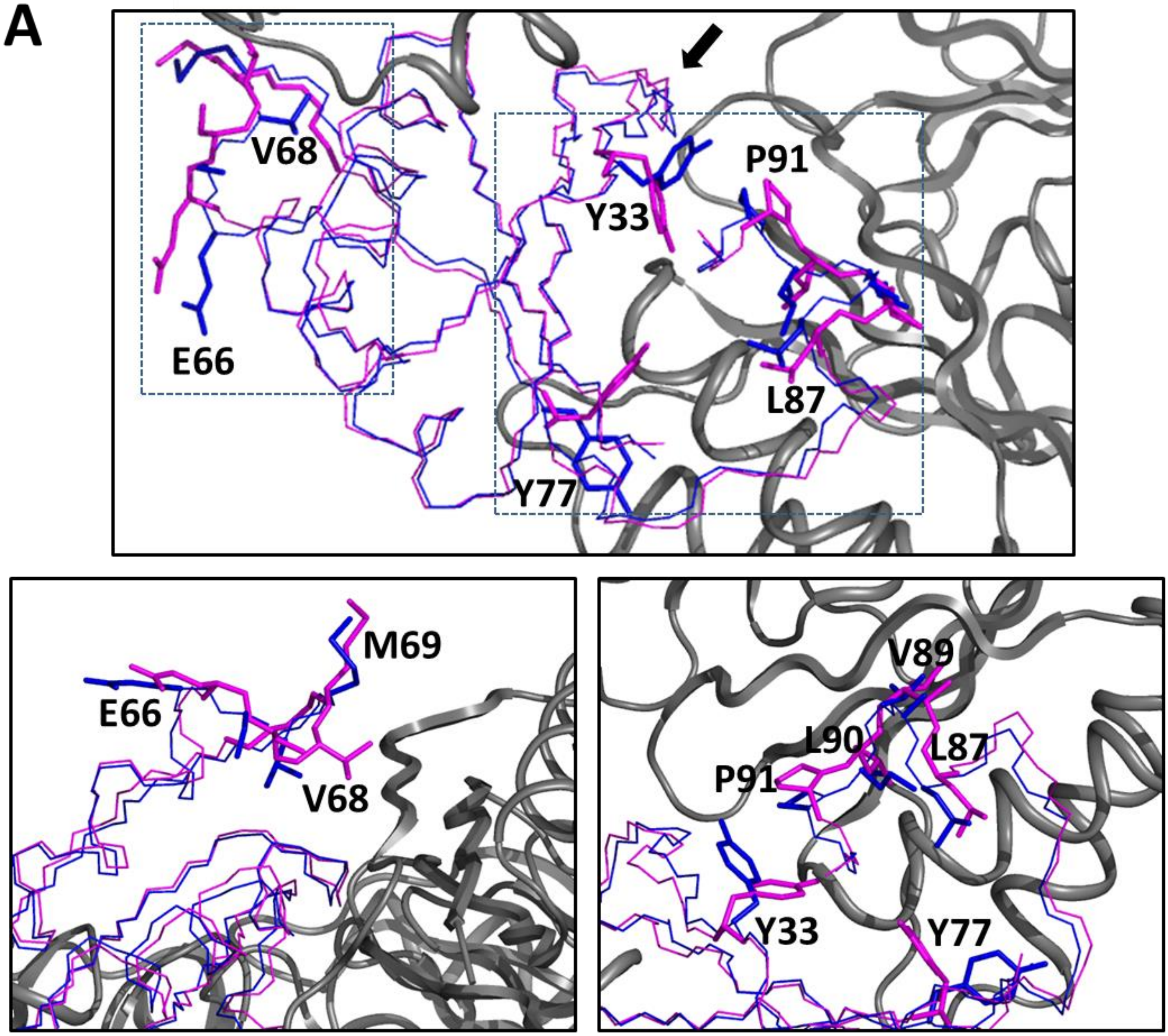

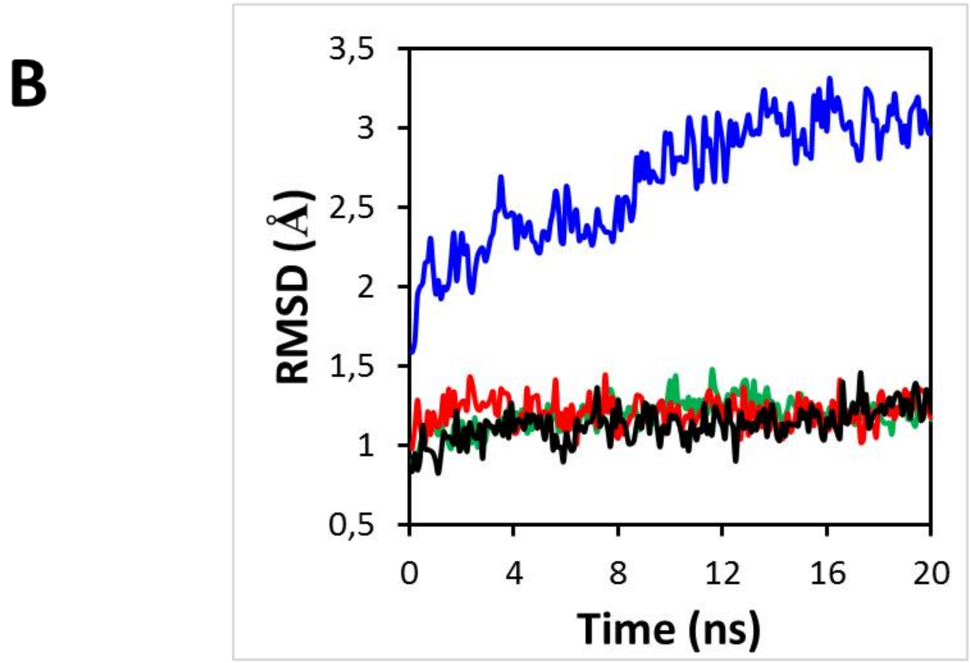

### Supplemental Figure 6

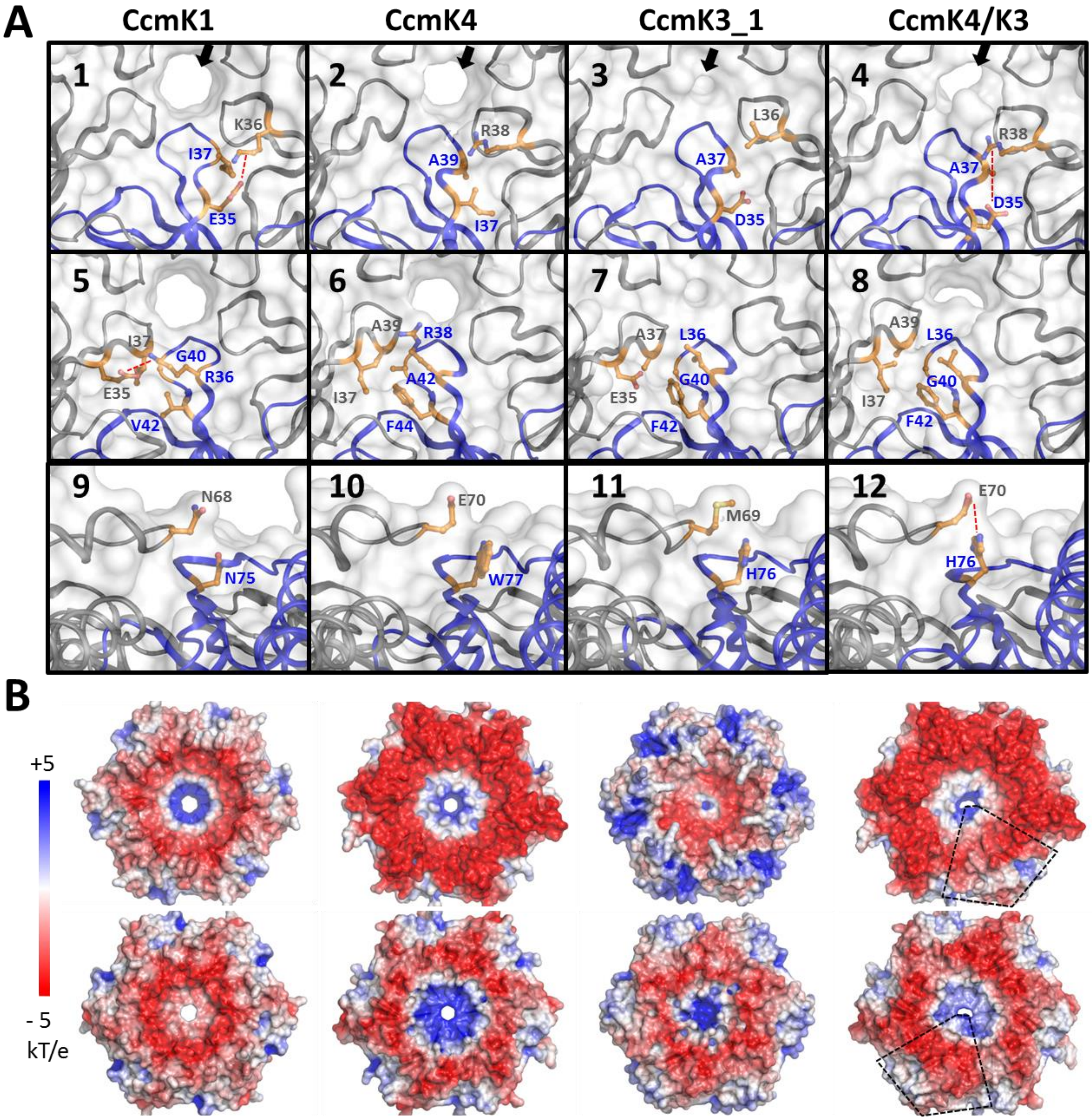

### Supplemental Figure 7

#### Figure S7

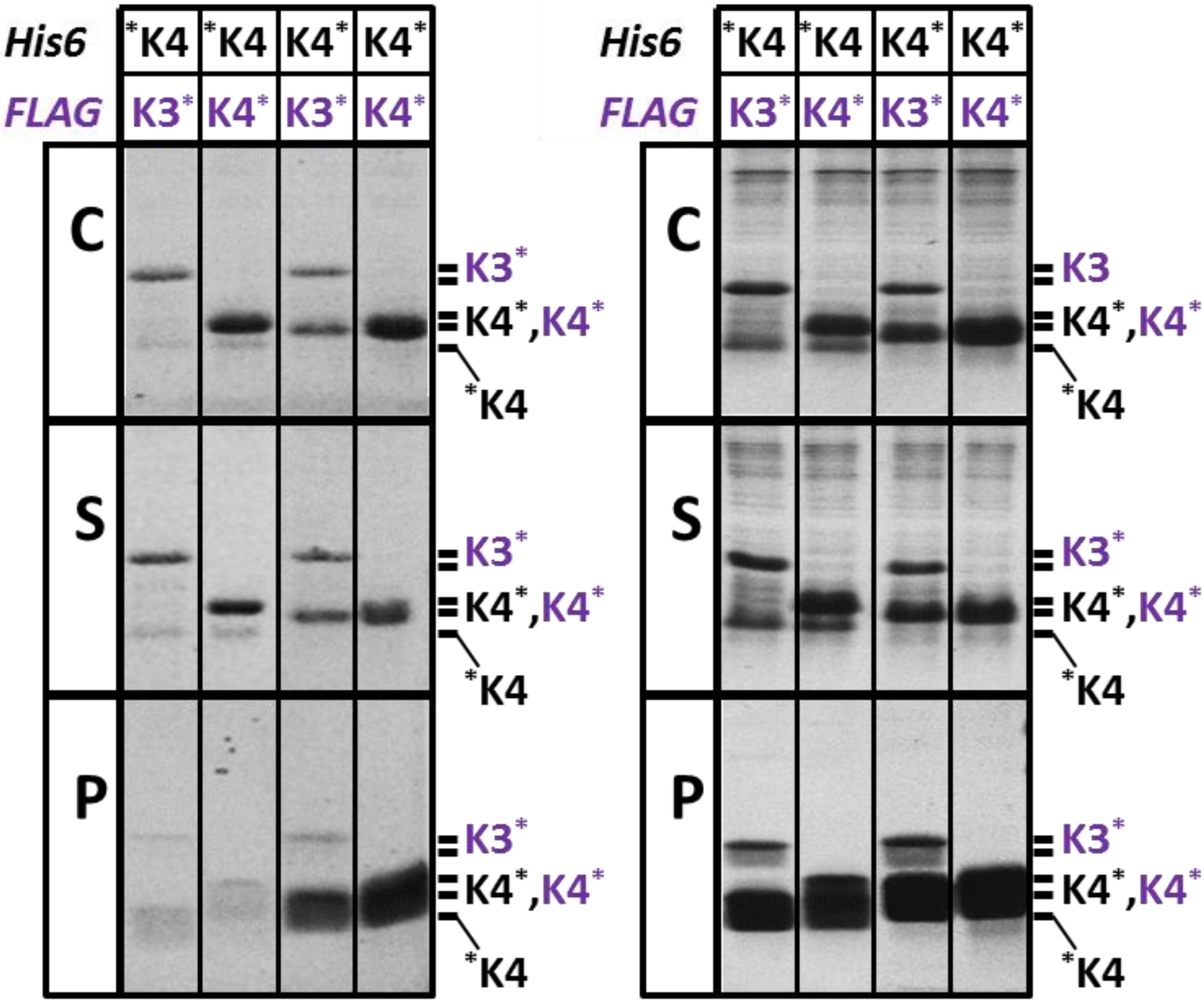
