## Supplemental List 1 for "Occurrence and stability of hetero-hexamer associations formed by β-carboxysome CcmK shell components"

**LIST S1 –** Preparation of pET26b-based vectors for studies of CcmK coexpression

The sequence indicated contains 4 cassettes flanked by T7 promoter/terminator sequences and containing identical RBS sequences.
Flanking SacI/KpnI sites permitted the integration in pBlueScript II SK+, for subsequent DNA manipulations. BglII and BlpI sites (underlined-italics) served for final transfer to pET26b.Other indicated sites permit the insertion of CcmK sequences (given below) carrying different tags at either N or C-ter. They are shown following the next order on the sequence: SwaI, BamHI, PacI, AgeI, MfeI, SalI, BsrGI, HindIII.

Studies of co-expression of CcmK couples were carried out using a two-cassette plasmid generated from the previous one as follows: 1) AvrII digestion and recircularization with T4 ligase (Thermofisher) to remove the 4$^{th}$ cassette; 2) BamHI and AgeI to remove the 2$^{nd}$ cassette, followed by treatment with the Klenow fragment LC (Thermofisher), and recircularization with T4 ligase.

GAGCTCTAA*AGATCT*CGATCCCGCGAAATTAATACGACTCACTATAGGGGAATTGTGAGCGGATAACAATTCCCCTCTAGAA<br>
ATAAGATTTAAATACTTTAAGAAGGAGATATACC**ATG**GCACATCACCATCATGCTAGCGGCGAAAATCTGTACTTCCAGGGT<br>
GCCATGGCAATCGAGAGTGCGGCCGCACATCACCATCATTGATAGGATCCACTTCTCGAGTTAACTCGTGAGCAATAACTAG<br>
CATAACCCCTTGGGGCCTCTAAACGGGTCTTGAGGGGTTTTTTGCTGAAAGTACACGGCCGCATAATCGAAATTAATACGAC<br>
TCACTATAGGGGAATTGTGAGCGGATAACAATTCCCCTCTAGAGTTAATTAAGTAAGTATAAGAAGGAGATATACC**ATG**GCA<br>
AGCTGGAGCCACCCGCAGTTCGAAAAGGGTGCTAGCGGCGAAAATCTGTACTTCCAGGGTGCCATGGCAATCGCTGTGTTCC<br>
AGAGTGCGGCCGCATGGAGCCACCCGCAGTTCGAAAAGTGACTAGTTACCGGTCACCTCTCGAGAGCAATAACTAGCATAAC<br>
CCCTTGGGGCCTCTAAACGGGTCTTGAGGGGTTTTTTGCTGAAAGTACACTCCAGCATTACGAAATTAATACGACTCACTAT<br>
AGGGGAATTGTGAGCGGATAACAATTCCCCTCTAGAAATAATTTTACAATTGTTTAAGAAGGAGATATACC**ATG**GATTACAA<br>
AGATGACGATGATAAGGCTAGCGGCGAAAATCTGTACTTCCAGGGTGCCATGGCACAAGCGGTGGAGTGCGGCCGCAGATTA<br>
CAAAGATGACGATGATAAGTGACTAGTATGTCGACTCCTAGGACTCGAGCAATAACTAGCATAACCCCTTGGGGCCTCTAAA<br>
CGGGTCTTGAGGGGTTTTTTGCTGAAACGATCCCGCGAAATTAATACGACTCACTATAGGGGAATTGTGAGCGGATAACAAT<br>
TCCCCTCTAGAAATAATGTACATTAACTTTAAGAAGGAGATATACC**ATG**GCATACCCGTACGATGTTCCGGATTACGCTAGC<br>
GGCGAAAATCTGTACTTCCAGGGTGCCATGGCAGCCCACCAGAGTGCGGCCGCATACCCGTACGATGTTCCGGATTACGCAT<br>
GACCTAGGAAAGCTTTCTCGAGA*GCTGAGC*AATAACTAGCATAACCCCTTGGGGCCTCTAAACGGGTCTTGAGGGGTTTTTT<br>
GGTACCAATTC

---

**SEQUENCES OF INDIVIDUAL GENES**

UNDERLINED: Flanking XbaI/XhoI sites are for transfer to PET15b and analysis of expression/solubility of individual protein constructs.
Underlined for His4-, StrepTag-, FLAG- and HA-tagged sequences are SwaI/BamHI, PacI/AgeI, MunI/SpeI and BsrGI/HindiIII sites required for integration at 1$^{st}$ to 4$^{th}$ cassette of the pET26-based vector for coexpression studies, respectively (highlighted above with cyan and yellow background colors).

Construct start and stop codons are indicated in blue bold, black bold letters being used for Ccm start codon within N-ter tagged sequences. Purification/labeling tags are shown in red, TEV-cleavage sequence in green.

> untagged CcmK1 Syn6803

CCTCTAGAAATAAGATTTAAATACTTTAAGAAGGAGATATACC**ATG**GCAATCGCTGTAGGTATGATCGAAACTCTGGGGT
TTCCGGCTGTTGTGGAAGCAGCCGATAGCATGGTAAAAGCGGCGCGCGTGACCTTAGTGGGCTATGAAAAGATTGGCAGC
GGTCGTGTCACCGTTATTGTTCGCGGGGATGTCAGCGAGGTGCAAGCGTCAGTGACGGCGGGTATCGAAAATATCCGTCG
TGTAAACGGTGGAGAAGTACTGTCAAACCATATCATCGCACGCCCACATGAAAATCTGGAGTATGTTTTACCGATTCGCT
ATACGGAAGCTGTGGAGCAGTTTCGTGAGATTGTAAACCCAAGCATCATCCGCCGTGCT**TAA**GATGTTTACTCCCGGGTA
GCGGCGAAAATCTGTACTTCCAGAGTGCGGCCGCACATCACCATCATTGATAGGATCCACTTCTCGAGGAT

> His4_TEV_CcmK1 Syn6803

CCTCTAGAAATAAGATTTAAATACTTTAAGAAGGAGATATACC**ATG**GCACATCACCATCATGCTAGCGGCGAAAATCTGT
ACTTCCAGGGTGCC**ATG**GCAATCGCTGTAGGTATGATCGAAACTCTGGGGTTTCCGGCTGTTGTGGAAGCAGCCGATAGC
ATGGTAAAAGCGGCGCGCGTGACCTTAGTGGGCTATGAAAAGATTGGCAGCGGTCGTGTCACCGTTATTGTTCGCGGGGA
TGTCAGCGAGGTGCAAGCGTCAGTGACGGCGGGTATCGAAAATATCCGTCGTGTAAACGGTGGAGAAGTACTGTCAAACC
ATATCATCGCACGCCCACATGAAAATCTGGAGTATGTTTTACCGATTCGCTATACGGAAGCTGTGGAGCAGTTTCGTGAG
ATTGTAAACCCAAGCATCATCCGCCGTGCT**TAA**GATGTTTACTCCCGGGTAGCGGCGAAAATCTGTACTTCCAGAGTGCG
GCCGCACATCACCATCATTGATAGGATCCACTTCTCGAGGAT

> Strep_TEV_CcmK1 Syn6803

CCTCTAGAGTTAATTAAGTAAGTATAAGAAGGAGATATACC**ATG**GCAAGCTGGAGCCACCCGCAGTTCGAAAAGGGTGCT
AGCGGCGAAAATCTGTACTTCCAGGGTGCC**ATG**GCAATCGCTGTAGGTATGATCGAAACTCTGGGGTTTCCGGCTGTTGT
GGAAGCAGCCGATAGCATGGTAAAAGCGGCGCGCGTGACCTTAGTGGGCTATGAAAAGATTGGCAGCGGTCGTGTCACCG
TTATTGTTCGCGGGGATGTCAGCGAGGTGCAAGCGTCAGTGACGGCGGGTATCGAAAATATCCGTCGTGTAAACGGTGGA
GAAGTACTGTCAAACCATATCATCGCACGCCCACATGAAAATCTGGAGTATGTTTTACCGATTCGCTATACGGAAGCTGT
GGAGCAGTTTCGTGAGATTGTAAACCCAAGCATCATCCGCCGTGCT**TAA**GATGTTTACTCCCGGGTAGCGGCGAAAATCT
GTACTTCCAGAGTGCGGCCGCATGGAGCCACCCGCAGTTCGAAAAGTGACTAGTTACCGGTCACCTCTCGAGGAT

> FLAG_TEV_CcmK1 Syn6803

CCTCTAGAAATAATTTTACAATTGTTTAAGAAGGAGATATACC**ATG**GATTACAAAGATGACGATGATAAGGCTAGCGGCG
AAAATCTGTACTTCCAGGGTGCC**ATG**GCAATCGCTGTAGGTATGATCGAAACTCTGGGGTTTCCGGCTGTTGTGGAAGCA
GCCGATAGCATGGTAAAAGCGGCGCGCGTGACCTTAGTGGGCTATGAAAAGATTGGCAGCGGTCGTGTCACCGTTATTGT
TCGCGGGGATGTCAGCGAGGTGCAAGCGTCAGTGACGGCGGGTATCGAAAATATCCGTCGTGTAAACGGTGGAGAAGTAC
TGTCAAACCATATCATCGCACGCCCACATGAAAATCTGGAGTATGTTTTACCGATTCGCTATACGGAAGCTGTGGAGCAG
TTTCGTGAGATTGTAAACCCAAGCATCATCCGCCGTGCT**TAA**GATGTTTACTCCCGGGTAGCGGCGAAAATCTGTACTTC
CAGAGTGCGGCCGCAGATTACAAAGATGACGATGATAAGTGACTAGTATGTCGACTCCTAGGACTCGAGGAT

> HA_TEV_CcmK1 Syn6803

CCTCTAGAAATAATGTACATTAACTTTAAGAAGGAGATATACC**ATG**GCATACCCGTACGATGTTCCGGATTACGCTAGCG
GCGAAAATCTGTACTTCCAGGGTGCC**ATG**GCAATCGCTGTAGGTATGATCGAAACTCTGGGGTTTCCGGCTGTTGTGGAA
GCAGCCGATAGCATGGTAAAAGCGGCGCGCGTGACCTTAGTGGGCTATGAAAAGATTGGCAGCGGTCGTGTCACCGTTAT
TGTTCGCGGGGATGTCAGCGAGGTGCAAGCGTCAGTGACGGCGGGTATCGAAAATATCCGTCGTGTAAACGGTGGAGAAG
TACTGTCAAACCATATCATCGCACGCCCACATGAAAATCTGGAGTATGTTTTACCGATTCGCTATACGGAAGCTGTGGAG
CAGTTTCGTGAGATTGTAAACCCAAGCATCATCCGCCGTGCT**TAA**GATGTTTACTCCCGGGTAGCGGCGAAAATCTGTAC
TTCCAGAGTGCGGCCGCATACCCGTACGATGTTCCGGATTACGCATGACCTAGGAAAGCTTTCTCGAGGAT

> CcmK1_TEV_His4 Syn6803

CCTCTAGAAATAAGATTTAAATACTTTAAGAAGGAGATATACC**ATG**GCAATCGCTGTAGGTATGATCGAAACTCTGGGGT
TTCCGGCTGTTGTGGAAGCAGCCGATAGCATGGTAAAAGCGGCGCGCGTGACCTTAGTGGGCTATGAAAAGATTGGCAGC
GGTCGTGTCACCGTTATTGTTCGCGGGGATGTCAGCGAGGTGCAAGCGTCAGTGACGGCGGGTATCGAAAATATCCGTCG
TGTAAACGGTGGAGAAGTACTGTCAAACCATATCATCGCACGCCCACATGAAAATCTGGAGTATGTTTTACCGATTCGCT
ATACGGAAGCTGTGGAGCAGTTTCGTGAGATTGTAAACCCAAGCATCATCCGCCGTGCGGGTAGCGGCGAAAATCTGTAC
TTCCAGAGTGCGGCCGCACATCACCATCAT**TGA**TAGGATCCCGGTCACCTCTCGAGAGCA

> CcmK1_TEV_Strep Syn6803

CCTCTAGAGTTAATTAAGTAAGTATAAGAAGGAGATATACC**ATG**GCAATCGCTGTAGGTATGATCGAAACTCTGGGGTTT
CCGGCTGTTGTGGAAGCAGCCGATAGCATGGTAAAAGCGGCGCGCGTGACCTTAGTGGGCTATGAAAAGATTGGCAGCGG
TCGTGTCACCGTTATTGTTCGCGGGGATGTCAGCGAGGTGCAAGCGTCAGTGACGGCGGGTATCGAAAATATCCGTCGTG
TAAACGGTGGAGAAGTACTGTCAAACCATATCATCGCACGCCCACATGAAAATCTGGAGTATGTTTTACCGATTCGCTAT
ACGGAAGCTGTGGAGCAGTTTCGTGAGATTGTAAACCCAAGCATCATCCGCCGTGCGGGTAGCGGCGAAAATCTGTACTT

CCAGAGTGCGGCCGCATGGAGCCACCCGCAGTTCGAAAAG**TGA**CTAGTTACCGGTCACCTCTCGAGGAT

> CcmK1_TEV_FLAG Syn6803
CCTCTAGAAATAATTTTACAATTGTTTAAGAAGGAGATATACC**ATG**GCAATCGCTGTAGGTATGATCGAAACTCTGGGGT
TTCCGGCTGTTGTGGAAGCAGCCGATAGCATGGTAAAAGCGGCGCGCGTGACCTTAGTGGGCTATGAAAAGATTGGCAGC
GGTCGTGTCACCGTTATTGTTCGCGGGGATGTCAGCGAGGTGCAAGCGTCAGTGACGGCGGGTATCGAAAATATCCGTCG
TGTAAACGGTGGAGAAGTACTGTCAAACCATATCATCGCACGCCCACATGAAAATCTGGAGTATGTTTTACCGATTCGCT
ATACGGAAGCTGTGGAGCAGTTTCGTGAGATTGTAAACCCAAGCATCATCCGCCGTGCGGGTAGCGGCGAAAATCTGTAC
TTCCAGAGTGCGGCCGCAGATTACAAAGATGACGATGATAAG**TGA**CTAGTATGTCGACTCCTAGGACTCGAGGAT

> CcmK1_TEV_HA Syn6803
CCTCTAGAAATAATGTACATTAACTTTAAGAAGGAGATATACC**ATG**GCAATCGCTGTAGGTATGATCGAAACTCTGGGGT
TTCCGGCTGTTGTGGAAGCAGCCGATAGCATGGTAAAAGCGGCGCGCGTGACCTTAGTGGGCTATGAAAAGATTGGCAGC
GGTCGTGTCACCGTTATTGTTCGCGGGGATGTCAGCGAGGTGCAAGCGTCAGTGACGGCGGGTATCGAAAATATCCGTCG
TGTAAACGGTGGAGAAGTACTGTCAAACCATATCATCGCACGCCCACATGAAAATCTGGAGTATGTTTTACCGATTCGCT
ATACGGAAGCTGTGGAGCAGTTTCGTGAGATTGTAAACCCAAGCATCATCCGCCGTGCGGGTAGCGGCGAAAATCTGTAC
TTCCAGAGTGCGGCCGCATACCCGTACGATGTTCCGGATTACGCA**TGA**CCTAGGAAAGCTTTCTCGAGGAT

All other CcmK Syn6803 sequences were mounted on the same way, just replacing CcmK1 above (blocks in grey above) by next DNA sequences:

> FOR CcmK2 Syn6803 constructs
GCAATCGCTGTGGGTATGATCGAAACACGCGGGTTTCCAGCGGTTGTGGAGGCGGCGGATTCAATGGTAAAAGCAGCGCGCG
TTACCTTAGTGGGCTATGAAAAGATTGGCAGCGGTCGTGTAACCGTTATTGTGCGTGGGGATGTTAGCGAAGTCCAGGCAAG
CGTCAGCGCCGGCATCGAGGCGGCAAATCGTGTGAATGGTGGGGAAGTACTGTCAACGCATATCATCGCACGCCCACATGAA
AATCTGGAGTATGTTTTACCGATCCGTTATACCGAAGAAGTTGAACAGTTCCGTACGTACG

> FOR CcmK3 Syn6803 constructs
GCACAAGCGGTGGGAGTGATTCAAACCTTGGGCTTTCCGAGCGTGTTAGCGGCGGCGGATGCGATGCTAAAAGGGGGCCGGG
TGACGCTGGTGTATTATGACCTGGCTGAACGAGGCAACTTTGTAGTAGCAATCCGAGGTCCCGTATCAGAGGTTAACCTTTC
GATGAAGATGGGATTAGCAGCGGTAAACGAGTCCGTCATGGGAGGTGAAATCGTTAGCCATTATATTGTGCCGAACCCGCCC
GAAAATGTGCTGGCGGTTCTGCCAGTGGAGTATACCGAAAAGGTTGCTCGTTTCCGGA

> FOR CcmK4 Syn6803 constructs
GCAGCCCAGAGCGCCGTGGGCAGCATTGAAACCATTGGCTTTCCGGGCATTCTTGCCGCCGCGGATGCGATGGTAAAAGCTG
GTCGCATTACCATTGTGGGCTATATTCGTGCGGGCTCTGCGCGCTTTACGCTGAACATTCGTGGGGATGTGCAGGAAGTTAA
AACGGCGATGGCTGCGGGCATCGATGCCATCAACCGTACAGAAGGAGCCGATGTGAAAACCTGGGTCATTATTCCGCGCCCA
CATGAAAATGTCGTTGCGGTTCTGCCGATCGATTTTAGCCCTGAAGTAGAACCCTTTCGCGAAGCAGCGGAGGGCCTGAACC
GTCGCG

---

All other individual cases or combinations implying CcmK from Syn7942 were mounted on the same way, just replacing in the corresponding construct the sequence spreading from the start to the stop codons (bold blue font).

> CcmK3_His6 Syn7942 CONSTRUCTS
**ATG**CCAATCGCAGTGGGAACCATCCAAACCTTGGGCTTCCCGCCCATCATCGCAGCGGCCGATGCAATGGTAAAAGCCGC
GCGGGTGACGATCACGCAGTACGGCCTCGCGGAGAGCGCACAGTTCTTTGTCAGCGTACGTGGGCCGGTCAGCGAGGTGG
AGACCGCAGTTGAGGCGGGGCTGAAAGCGGTGGCGGAGACCGAGGGCGCCGAATTGATTAACTATATTGTGATCCCAAAC
CCACAGGAGAACGTGGAAACTGTGATGCCCATTGACTTCACGGCAGAATCAGAACCCTTTCGAAGCGGTAGTGCGGCCGC
AGGTAGTGGCGGTGCACATCACCATCACCATCAT**TGA**

> CcmK3_FLAG Syn7942 CONSTRUCTS
**ATG**CCAATCGCAGTGGGAACCATCCAAACCTTGGGCTTCCCGCCCATCATCGCAGCGGCCGATGCAATGGTAAAAGCCGC
GCGGGTGACGATCACGCAGTACGGCCTCGCGGAGAGCGCACAGTTCTTTGTCAGCGTACGTGGGCCGGTCAGCGAGGTGG
AGACCGCGGTGGAGGCGGGGCTGAAGGCGGTGGCGGAGACCGAGGGCGCCGAATTGATTAACTATATTGTGATCCCAAAC
CCACAGGAGAACGTGGAAACTGTGATGCCCATTGACTTCACGGCAGAATCAGAACCCTTTCGAAGCGGGAGTGCGGCCGC
AGGTAGTGGCGGTGCAGATTACAAAGATGACGATGATAAG**TGA**

> His6_CcmK4 Syn7942 CONSTRUCTS
**ATG**GCACATCACCATCACCATCATGCTAGCGGCGGTTCTGGTGGCATGTCACAGCAAGCGATTGGCAGCCTGGAGACCAA
GGGCTTTCCGCCCATTCTCGCCGCCGCCGATGCAATGGTAAAAGCTGGTCGTATCACCATCGTGAGCTATATGCGGGCGG
GTAGCGCCCGCTTTGCGGTTAACATCCGGGGCGACGTGTCTGAAGTCAAAACTGCGATGGACGCGGGTATCGAAGCTGCT
AAAAATACCCCAGGTGGTACGCTGGAGACCTGGGTTATTATCCCGCGACCGCACGAAAATGTGGAAGCCGTGTTTCCAAT
CGGTTTTGGCCCAGAAGTAGAACAATACCGGCTGTCAGCGGAGGGCACTGGCTCGGGTCGTCGC**TGA**

> CcmK4_His6 Syn7942 CONSTRUCTS
**ATG**TCACAGCAAGCGATTGGCAGCCTGGAGACCAAGGGCTTTCCGCCCATTCTCGCCGCCGCCGATGCAATGGTAAAAGC
TGGTCGTATCACCATCGTGAGCTATATGCGGGCGGGTAGCGCCCGCTTTGCGGTTAACATCCGGGGCGACGTGTCTGAAG
TCAAAACTGCGATGGACGCGGGTATCGAAGCTGCTAAAAATACCCCAGGTGGTACGCTGGAGACCTGGGTTATTATCCCG
CGACCGCACGAAAATGTGGAAGCCGTGTTTCCAATCGGTTTTGGCCCAGAAGTAGAACAATACCGGCTGTCAGCGGAGGG
CACTGGCTCGGGTCGTCGCGGTAGTGCGGCCGCAGGTAGTGGCGGTGCACATCACCATCACCATCAT**TGA**

> CcmK4_FLAG Syn7942 CONSTRUCTS
**ATG**TCACAGCAAGCGATTGGCAGCCTGGAGACCAAGGGCTTTCCGCCCATTCTCGCCGCCGCCGATGCAATGGTAAAAGC
TGGTCGTATCACCATCGTGAGCTATATGCGGGCGGGTAGCGCCCGCTTTGCGGTTAACATCCGGGGCGACGTGTCTGAAG
TCAAAACTGCGATGGACGCGGGTATCGAAGCTGCTAAAAATACCCCAGGTGGTACGCTGGAGACCTGGGTTATTATCCCG
CGACCGCACGAAAATGTGGAAGCCGTGTTTCCAATCGGTTTTGGCCCAGAAGTAGAACAATACCGGCTGTCAGCGGAGGG
CACTGGCTCGGGTCGTCGCGGTAGTGCGGCCGCAGGTAGTGGCGGTGCAGATTACAAAGATGACGATGATAAG**TGA**
