## Supplemental Table 1 for "Occurrence and stability of hetero-hexamer associations formed by β-carboxysome CcmK shell components"

**Table S1 – Analysis of native-MS and MS-MS data collected on CcmK1/K2 and CcmK3/K4 hetero-hexamers.**

| Sample | Theor. MW[a] | Exp. MW | Stoich. | Deviation[b] |
|---|---|---|---|---|
| **K1*/K2*** | 13153.0 (K1m) | 12988 | 1 | - 165 |
| | 12185.8 (K2m) | 12041 | 1 | -145 |
| | 73114.8 (K2h) | | 6 | |
| | | 73184 | 1:5 | -9 |
| | | **74212** | 2:4 | +72 |
| | | **75143** | 3:3 | +56 |
| | | 76078 | 4:2 | +44 |
| | 78918.0 (K1h) | | | |
| | | 122346 | 2:8 | +42 |
| | | 123380 | 3:7 | +129 |
| | | 124206 | 4:6 | +8 |
| **K1* + K2*** | 13153.0 (K1m) | 13012 | 1 | -141 |
| | 78918.0 (K1h) | 77694 | 6:0 | -234 |
| | | **78123** | 6:0 | +195 |
| ***K4/K3*** | 12085.8 (K4m) | 12087 | 1 | +1 |
| | 11972.8 (K3m) | 11842 | 1 | -131 |
| | | 12094 | 1 | +8 (K4) |
| | 71836.8 (K3h) | | | |
| | | 37783 | NA | |
| | | 66852 | NA | |
| | | 71087 | 0:6 | +35 |
| | | **72334** | 5:1 | +57 |
| | 72514.8 (K4h) | 72568 | 6:0 | +46 |

[a] Theoretical molecular weights calculated from amino acid sequences. The values of theoretical hexamers are also given (indicated by h letter following the isoform name). [b] Mass deviations between experimental and theoretical values are indicated. When considering monomers, the deviation is with regard to theoretical molecular weight; In the case of mixed associations, the deviation is with regard to values calculated for such combinations of experimental masses measured for each component as monomer. This is basically to account for typical losses of first methionine residue (131.04 Da) with components tagged at C-terminus.
