## Supplemental Table 2 for "Occurrence and stability of hetero-hexamer associations formed by β-carboxysome CcmK shell components"

### Table S2A – CcmK sequence identity scores considering full sequences

| % | K1_6803 | K1_7418 | K2_6803 | K2_7942 | K2_7418 | K3_6803 | K3_7942 | K3_7418 | K4_6803 | K4_7942 | K4_7418 |
|---|---|---|---|---|---|---|---|---|---|---|---|
| K1_6803 | 100,0 | 83,8 | 91,3 | 89,2 | 90,1 | 48,0 | 48,0 | 49,0 | 49,5 | 46,9 | 51,4 |
| K1_7418 | 83,8 | 100,0 | 88,4 | 89,2 | 92,1 | 49,0 | 48,0 | 51,0 | 48,2 | 46,4 | 52,2 |
| K2_6803 | 91,3 | 88,4 | 100,0 | 89,2 | 92,1 | 50,0 | 49,0 | 52,0 | 53,4 | 51,5 | 55,3 |
| K2_7942 | 89,2 | 89,2 | 89,2 | 100,0 | 95,1 | 52,0 | 51,0 | 52,0 | 52,0 | 50,0 | 54,9 |
| K2_7418 | 90,1 | 92,1 | 92,1 | 95,1 | 100,0 | 51,5 | 49,5 | 51,5 | 52,5 | 48,5 | 54,5 |
| K3_6803 | 48,0 | 49,0 | 50,0 | 52,0 | 51,5 | 100,0 | 52,0 | 58,8 | 46,1 | 45,1 | 51,0 |
| K3_7942 | 48,0 | 48,0 | 49,0 | 51,0 | 49,5 | 52,0 | 100,0 | 60,8 | 54,9 | 52,9 | 50,0 |
| K3_7418 | 49,0 | 51,0 | 52,0 | 52,0 | 51,5 | 58,8 | 60,8 | 100,0 | 50,0 | 51,0 | 51,0 |
| K4_6803 | 49,5 | 48,2 | 53,4 | 52,0 | 52,5 | 46,1 | 54,9 | 50,0 | 100,0 | 69,4 | 65,8 |
| K4_7942 | 46,9 | 46,4 | 51,5 | 50,0 | 48,5 | 45,1 | 52,9 | 51,0 | 69,4 | 100,0 | 66,4 |
| K4_7418 | 51,4 | 52,2 | 55,3 | 54,9 | 54,5 | 51,0 | 50,0 | 51,0 | 65,8 | 66,4 | 100,0 |

### Table S2B – CcmK sequence identity scores considering only inter-monomer residues

| % | K1_6803 | K1_7418 | K2_6803 | K2_7942 | K2_7418 | K3_6803 | K3_7942 | K3_7418 | K4_6803 | K4_7942 | K4_7418 |
|---|---|---|---|---|---|---|---|---|---|---|---|
| K1_6803 | 100,0 | 95,5 | 95,5 | 97,7 | 95,5 | 47,7 | 45,5 | 43,2 | 56,8 | 50,0 | 63,6 |
| K1_7418 | 95,5 | 100,0 | 95,5 | 95,5 | 100,0 | 47,7 | 43,2 | 43,2 | 54,6 | 47,7 | 61,4 |
| K2_6803 | 95,5 | 95,5 | 100,0 | 95,5 | 95,5 | 45,5 | 43,2 | 43,2 | 56,8 | 50,0 | 63,6 |
| K2_7942 | 97,7 | 95,5 | 95,5 | 100,0 | 95,5 | 50,0 | 45,5 | 45,5 | 56,8 | 50,0 | 63,6 |
| K2_7418 | 95,5 | 100,0 | 95,5 | 95,5 | 100,0 | 47,7 | 43,2 | 43,2 | 54,6 | 47,7 | 61,4 |
| K3_6803 | 47,7 | 47,7 | 45,5 | 50,0 | 47,7 | 100,0 | 52,3 | 65,9 | 45,5 | 43,2 | 50,0 |
| K3_7942 | 45,5 | 43,2 | 43,2 | 45,5 | 43,2 | 52,3 | 100,0 | 61,4 | 59,1 | 59,1 | 52,3 |
| K3_7418 | 43,2 | 43,2 | 43,2 | 45,5 | 43,2 | 65,9 | 61,4 | 100,0 | 52,3 | 54,6 | 54,6 |
| K4_6803 | 56,8 | 54,6 | 56,8 | 56,8 | 54,6 | 45,5 | 59,1 | 52,3 | 100,0 | 81,8 | 84,1 |
| K4_7942 | 50,0 | 47,7 | 50,0 | 50,0 | 47,7 | 43,2 | 59,1 | 54,6 | 81,8 | 100,0 | 77,3 |
| K4_7418 | 63,6 | 61,4 | 63,6 | 63,6 | 61,4 | 50,0 | 52,3 | 54,6 | 84,1 | 77,3 | 100,0 |

### Table S2C – Average CcmK sequence identities scores

<table>
<tr><th>%</th><th>K1_6803</th><th>K1_7418</th><th>K2_6803</th><th>K2_7942</th><th>K2_7418</th><th>K3_6803</th><th>K3_7942</th><th>K3_7418</th><th>K4_6803</th><th>K4_7942</th><th>K4_7418</th></tr>
<tr><th>K1_6803</th><td colspan="5" rowspan="5">90% full sequences<br>96% inter-monomer</td><td colspan="3" rowspan="5">50% full sequences<br>45% inter-monomer</td><td colspan="3" rowspan="5">51% full sequences<br>56% inter-monomer</td></tr>
<tr><th>K1_7418</th></tr>
<tr><th>K2_6803</th></tr>
<tr><th>K2_7942</th></tr>
<tr><th>K2_7418</th></tr>
<tr><th>K3_6803</th><td colspan="5" rowspan="3">50% full sequences<br>45% inter-monomer</td><td colspan="3" rowspan="3">56% full sequences<br>58% inter-monomer</td><td colspan="3" rowspan="3">50% full sequences<br>52% inter-monomer</td></tr>
<tr><th>K3_7942</th></tr>
<tr><th>K3_7418</th></tr>
<tr><th>K4_6803</th><td colspan="5" rowspan="3">51% full sequences<br>56% inter-monomer</td><td colspan="3" rowspan="3">50% full sequences<br>52% inter-monomer</td><td colspan="3" rowspan="3">68% full sequences<br>81% inter-monomer</td></tr>
<tr><th>K4_7942</th></tr>
<tr><th>K4_7418</th></tr>
</table>
