## Supplemental Table 3 for "Occurrence and stability of hetero-hexamer associations formed by β-carboxysome CcmK shell components"

**Table S3 – Crystallographic data and refinement statistics**

| | | |
|---|---|---|
| Cell parameters (Å, °) | $a = b = 70.68$, $c = 64.79$, $\alpha = \beta = 90$, $\gamma = 120$ | |
| Space group | $P6_3$ | |
| Resolution range (Å) | 61.21 – 1.80 | 1.84 – 1.80 |
| Nb. of observations | 178,945 | 10,458 |
| Nb. of unique reflections | 17,065 | 994 |
| Multiplicity | 10.5 | 10.5 |
| Completeness (%) | 99.6 | 99.7 |
| Rmerge | 0.095 | 1.341 |
| Rmeas | 0.100 | 1.348 |
| Rpim | 0.031 | 0.414 |
| I/σ | 14.1 | 1.9 |
| CC(1/2) | 0.999 | 0.847 |
| Resolution range (Å) | 61.20 – 1.80 | |
| Nb. of reflections | 17,024 | |
| Completeness | 99.4 | |
| Nb. of atoms | 1627 | |
| Protein | 1529 | |
| Water, ethylene glycol | 98 | |
| Rfactor | 0.1710 | |
| Rfree | 0.1924 | |
| rmsd bond lengths (Å) | 0.006 | |
| rmsd bond angles (°) | 0.733 | |
