## Supplemental Table 4 for "Occurrence and stability of hetero-hexamer associations formed by β-carboxysome CcmK shell components"

**Table S4 – Differential Scanning Fluorimetry and Dynamic Light Scattering results**

| | DSF | | | DLS | | | | |
|---|---|---|---|---|---|---|---|---|
| **Sample** | **$T_m$ (°C)** | **C.I. [a] (°C)** | **N.M.[b]** | **$T_{aggr}$ ( °C)** | **C.I. [a] (°C)** | **N.M.[b]** | **$R_h$ [c] (nm)** | **Pd [d] (%)** |
| $_{His4}$-K4 | 89,2[e] | 88,6 – 89,7 | 2 | NP[f] | | 2 | 4.2 | 16.0 |
| $_{His4}$-K4/K3-$_{FLAG}$ | 59,6 | 59,4 – 59,8 | 3 | 61,3 | 61,1 – 61,4 | 3 | 5.7 | 25.7 |
| K1-$_{His4}$/K1-$_{FLAG}$ | NP[f] | | 3 | NP[f] | | 3 | 5.2 | 29.0 |
| K2-$_{His4}$/K2-$_{FLAG}$ | 62,3[e] | 61,7 – 62,8 | 3 | 80,6 | 80,5 – 80,7 | 4 | 5.7 | 27.2 |
| K1-$_{His4}$/K2-$_{FLAG}$ | 58, 7 | 58,5 – 58,9 | 3 | 71,3 | 71,2 – 71,4 | 3 | 5.7 | 31.5 |

[a] 95% confidence interval. [b] Number of measurements. [c] Hydrodynamic radius. [d] Polydispersity index. [e] Calculated from a low fluorescence signal event. [f] No measurable transition observed.
