## Supplementary material for "Occurrence and stability of hetero-hexamer associations formed by β-carboxysome CcmK shell components": Legends for Supplementary Figures and Tables

Short title: Hetero-oligomerization of carboxysome shell components

Garcia-Alles Luis F.[1,*], Root Katharina[2], Maveyraud Laurent[3], Aubry Nathalie[1], Lesniewska Eric[4], Mourey Lionel[3], Zenobi Renato[2], Truan Gilles[1]

**Table S1 – Analysis of native-MS and MS-MS data collected on CcmK1/K2 and CcmK3/K4 hetero-hexamers.**

**Table S2 – CcmK sequence identity scores.**

**Table S3 – Crystallographic data and refinement statistics.**

**Table S4 – Differential Scanning Fluorimetry and Dynamic Light Scattering results.**

**List S1 – Preparation of pET26b-based vectors for studies of CcmK coexpression.**

**Figure S1 – Expression and solubility of untagged *Syn6803* CcmK paralogs in *Escherichia coli*.** Left panel: total cellular contents, right: material remaining soluble in supernatants after lysis and centrifugation at 20000 g. CcmK monomer bands are expected within the 10-15 kDa range. "Empty" corresponds to cells transformed with the negative control pET15b vector lacking the *ccmK* gene.

**Figure S2 – Expression and solubility of tagged *Syn6803* CcmK paralogs in *Escherichia coli*.** Coomassie-stained SDS-PAGE views of cellular and soluble fractions prepared from BL21(DE3) strains transformed with pET15b-based plasmids permitting expression of CcmK proteins tagged with the indicated peptides. On top, total cellular expression levels are shown, bottom part is for material remaining in supernatants after lysis and centrifugation at 20.000 g. *A*, results collected for the N-ter tagged protein versions. *B*, similar data collected for C-ter tagged proteins. Please notice that the relative vertical positioning of bands might differ slightly, as a consequence of the image mounting process.

**Figure S3 – Hetero-hexamer formation with combined *Syn6803* CcmK paralogs**. *A*, Schematic representation of constructed expression vectors. Two ORFs coding for CcmK proteins tagged at either N- or C-terminus with His4 and FLAG peptides were engineered between T7 promoter and terminator sequences (blue and empty circles, respectively). The combinations studied are listed below, in the table. Whether asterisk is written before or after the paralog name denotes N- or C-tagging emplacement, respectively. Black letters are used for His4-carrying subunit, violet for the FLAG-tagged proteins. *B*, Coomassie-stained SDS-PAGE showing total cellular contents (C), soluble material remaining in supernatants after lysis and centrifugation (S) and purified fractions (P). Only the portion of the gels presenting the region where CcmK monomers appear is shown. The approximate position of the different co-expressed partners is indicated on the right. Please notice that, as a consequence of partial proteolysis of C-ter CcmK4 proteins, their emplacement cannot be unambiguously indicated. Besides, the relative vertical positioning of bands in gels might be slightly erroneous, as a consequence of image mounting process. White arrows are to indicate the absence of CcmK1-$His_4$ soluble bands in combinations showing expression, whereas black arrows highlight CcmK1-FLAG and CcmK2-FLAG solubility that seems unaffected by the presence of CcmK3.

**Figure S4 – CcmK hetero-association occurrence upon co-expression of all *Syn6803* CcmK**. *A*, Schematic representation of expression vectors constructed. Four ORFs coding for CcmK proteins tagged at either N- or C-terminus with His4 (black), Strep (blue), FLAG (violet) or HA (red) peptides were engineered between T7 promoter and terminator sequences (blue and empty circles, respectively). The studied combinations are listed in the table below. Whether asterisk is written on before or after the paralog name denotes N- or C-tagging emplacement, respectively. *B*, Coomassie-stained SDS-PAGE showing total cellular contents. (C), soluble material remaining in supernatants after lysis and centrifugation (S) and purified fractions (P) prepared after culturing BL21(DE3) transformed with plasmids schematized in panel A in auto-induction media. Results are organized depending on the position of the FLAG-tag with regard to the protein placed on the third cassette. Only the portion of the gels presenting the region where CcmK monomers appear is shown. White arrows indicate an absence of CcmK1-$His_4$ soluble bands in combinations showing expression. *C*, Western blot analysis of TALON-purified fractions. Detection on PVDC membranes was effected stepwise, a first incubation being performed with a mixture of mouse antiFLAG and mouse antiHA, followed by a second incubation with a mixture containing a secondary IgG antimouse-AP fusion plus streptactin-AP conjugate. The vertical position of protein bands in images B and C are not strictly matched with each other. Please refer to M&M for further details.

**Figure S5 – Homology models of Syn*6803* CcmK3 and molecular dynamics**. *A,* Structures built by SWISS-MODEL (CcmK3_1, blue) and PHYRE2 (CcmK3_2, magenta), shown after alignment of the two colored monomers. For clarity, only the backbone trace is shown. Other neighbor monomers of the hexamer are represented as grey ribbons (only for CcmK3_1). Residues cumulating major differences are highlighted by thicker lines and include side-chain atoms. The two regions squared on the top view are zoomed and rotated in bottom images. The arrow indicates the position of residues lining the hypothetical pore, which would be clogged in CcmK3. *B*, Results of all-atom MD simulations run on CcmK3 homo-hexamer models and on CcmK3/K4 hetero-hexamers built after replacement of a single monomer of CcmK4 hexamer (PDB code 2A10) by either CcmK3_1 or CcmK3_2 models. RMSD deviations were measured for main-chain atoms of snapshot structures (recorded every 250 ps) compared to their position in the starting structure. Values estimated for the CcmK3_1 hexamer are shown in green, for CcmK3_2 hexamer in blue, CcmK3_1/CcmK4 hetero-hexamer in black and CcmK3_2/CcmK4 in red. Two independent 20 ns MD were run for each case, with analogous results.

**Figure S6 – Potential consequences on the structure and electrostatic properties of the formation of Syn*6803* CcmK3/K4 hetero-hexamers**. *A,* Comparison of structural details in different CcmK homo-hexamers, including the CcmK3_1 homology model built by SWISS-MODEL (CcmK3_1, blue), and the CcmK3_1/K4 hetero-hexamer model. All structures were pre-aligned to generate equivalent views. One of the monomers is shown in blue (corresponding to the CcmK3_1 subunit in the hetero-hexamer), neighbor ones are shown in grey ribbon representation. For clarity, only the backbone trace is shown. Residues that could contribute differently to inter-monomer contacts are colored as in Fig. 4, with side-chain atoms drawn as balls-and-sticks. Arrows indicate the position of central hexamer pores, which is clogged in CcmK3. Ionic interactions are illustrated as red dashes. *B*, Comparison of surfaces of homo-hexamers and the CcmK3/K4 hetero-hexamer, colored according to electrostatic potential. Top and bottom images correspond to views of concave and convex sides, respectively. The approximate emplacement of the CcmK3 subunit in the CcmK3/K4 heterohexamer is indicated with dashed lines.

**Figure S7 – Hetero-hexamer formation between *Syn7942* CcmK3 and CcmK4 proteins**. Coomassie-stained SDS-PAGE showing total cellular contents (C), soluble material remaining in supernatants after lysis and centrifugation (S), and TALON-purified fractions (P). Left panel correspond to samples prepared from cellular cultures that were induced in IPTG (200 µM), whereas on the right are presented data from samples produced under auto-induction conditions. Top lines indicate the combination of isoforms co-expressed together, as well as the tagging identity. Tag emplacement at either N-ter or C-ter is indicated with an asterisk written before or after the paralog abbreviation, respectively.
